# Flavonols and dihydroflavonols inhibit the main protease activity of SARS-CoV-2 and the replication of human coronavirus 229E

**DOI:** 10.1101/2021.07.01.450756

**Authors:** Yue Zhu, Frank Scholle, Samantha C. Kisthardt, De-Yu Xie

## Abstract

Since December 2019, the deadly novel severe acute respiratory syndrome coronavirus 2 (SARS-CoV-2) has caused the current COVID-19 pandemic. To date, vaccines are available in the developed countries to prevent the infection of this virus, however, medicines are necessary to help control COVID-19. Human coronavirus 229E (HCoV-229E) causes the common cold. The main protease (M^pro^) is an essential enzyme required for the multiplication of these two viruses in the host cells, and thus is an appropriate candidate to screen potential medicinal compounds. Flavonols and dihydroflavonols are two groups of plant flavonoids. In this study, we report docking simulation with two M^pro^ enzymes and five flavonols and three dihydroflavonols, *in vitro* inhibition of the SARS-CoV-2 M^pro^, and *in vitro* inhibition of the HCoV 229E replication. The docking simulation results predicted that (+)-dihydrokaempferol, (+)-dihydroquercetin, (+)-dihydromyricetin, kaempferol, quercetin, myricentin, isoquercetin, and rutin could bind to at least two subsites (S1, S1’, S2, and S4) in the binding pocket and inhibit the activity of SARS-CoV-2 M^pro^. Their affinity scores ranged from −8.8 to −7.4. Likewise, these compounds were predicted to bind and inhibit the HCoV-229E M^pro^ activity with affinity scores ranging from −7.1 to −7.8. *In vitro* inhibition assays showed that seven available compounds effectively inhibited the SARS-CoV-2 M^pro^ activity and their IC50 values ranged from 0.125 to 12.9 µM. Five compounds inhibited the replication of HCoV-229E in Huh-7 cells. These findings indicate that these antioxidative flavonols and dihydroflavonols are promising candidates for curbing the two viruses.

## 1. Introduction

SARS-CoV-2 is the abbreviation of the novel severe acute respiratory syndrome coronavirus 2. This virus was firstly reported to cause a severe pneumonia in December of 2019 in Wuhan, China [1–3]. On February 11, 2020, the World Health Organization (WHO) designated this pneumonia as coronavirus disease 2019 (COVID-19). COVID-19 the rapidly spread different countries. On March 11, 2020, WHO announced the COVID-19 pandemic [4, 5]. This pandemic has rapidly spread across all over the world. By June 21, 2021, based on the COVID-19 Dashboard by Center for Systems Science and Engineering at Johns Hopkins Coronavirus Resource Center, 117,553,726 infected cases and 3,867,641 deaths have been reported from more than 200 countries or regions. No strategy to stop the spread of this virus was available until January 2021, when several vaccines started to be approved for vaccination in several countries [6–11]. On one hand, since the start of vaccination, the number of infections has started to decrease. On the other hand, due to the insufficient vaccine quantities and vaccine hesitancy even where available, in the first week of February, 2021, the daily infection cases and deaths were still more than 400,000 and 11,000, respectively. By Feb. 9, the case numbers still increased by more than 300,000 daily. Meanwhile, the use of vaccines has also indicated that developing effective medicines is necessary to stop COVID-19. A recent study showed that mutations in the spike protein of SARS-CoV-2 might cause the escape of new variants from antibody [12]. The variant B.1.351 found in South Africa was reported to be able to escape vaccines developed by AstraZeneca, Johnson & Johnson (J&J), and Novavax [13]. Merck & Co has stopped their race for vaccines due to the lack of effectiveness of their products, instead, they continue to focus on antiviral drug development [14]. Unfortunately, to date, effective medicines are still under screening. Although chloroquine and hydroxychloroquine were reported to be potentially effective in helping to improve COVID-19 [15], the use of these two anti-malarial medicines has been arguable in USA because of potential risk concerns [16]. Other potential candidate medicines are the combination of α-interferon and anti-HIV drugs lopinavir/ritonavir [17], and remdesivir [15, 18]. Given that the efficacy of all these medicines being repurposed has not been conclusive, further studies are necessary to apply them for treating COVID-19.

SARS-CoV-2 is a single stranded RNA virus. Its genomic RNA contains around 30,000 nucleotides and forms a positive sense strand with a 5′ methylated cap and a 3′ polyadenylated tail that encodes at least six open reading frames (ORF) [19, 20]. This feature allows it to be able to use the ribosomes of the host cells to translate proteins. The longest ORF (ORF1a/b) translates two polyproteins, which are cleaved by one main protease (M^pro^, a 3C-like protease, 3CL^pro^) and another papain-like protease (PL2pro) into 16 nonstructural proteins (NSPs), which include RNA-dependent RNA polymerase (RdRp, nsp12), RNA helicase (nsp13), and exoribonuclease (nsp14). The NSPs subsequently produce structural and accessory proteins. The structural proteins include an envelope protein, membrane protein, spike (S) protein, and nucleocapsid protein (Fig. 1) [21, 22]. The S protein is a type of glycoprotein and plays an essential role in the attachment and the infection of the host cells [23]. It binds to the human angiotensin converting enzyme 2 to help the virus enter the human cells [24, 25]. Since May 2020, the mutations of amino acids of the S protein has created a large number of variants, of which the emergence of alpha, beta, gamma, and delta variants has shown more pathogenic and transmissible, thus caused potential challenges to use vaccines to completely control the pandemic [26–30].

**Figure 1.**
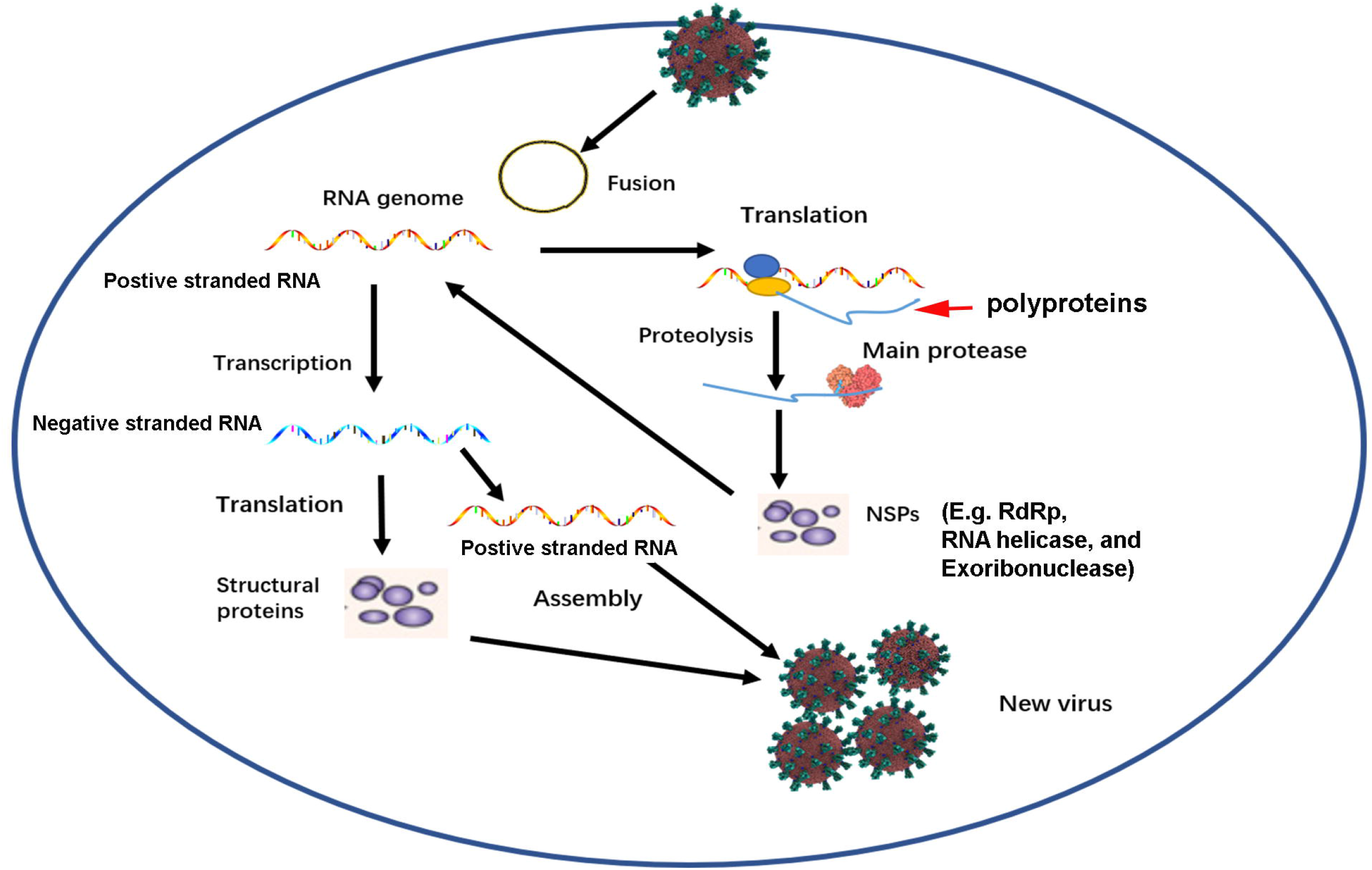
A diagram showing the function of the SARS-CoV-2 main protease in the virus replication in the host cells. Once the virus enters into the host cells. Its positive sense and single stranded RNA uses the ribosomes to translate open reading frames 1a and 1b to polyproteins (PP), in which the main protease and papain-like protease cleaves PPs to non-structural proteins (NSPs). Three NSPs, RNA dependent RNA polymerase (RdRp), RNA helicase, and exoribonuclease, are involved in the transcription of the positive RNA to negative sense and single stranded RNA, which is further transcribed to positive sense and single stranded RNA. Finally, structural proteins and a positive single stranded RNA assembly together to form a virus progeny.

Human coronavirus 229E (HCoV-229E) is a pathogenic virus in the genus *Alphacoronavirus* [31]. It is one of the causative viral agents of the common cold [32, 33]. Its genome consists of a positive sense and single-stranded RNA with 27,317 nucleotides (nt). Its genome size commonly varies in different clinical isolates. For example, HCoV-229E strains 0349 and J0304 were two clinical isolates causing the common cold [34]. The entire genome of these two clinical isolates were reported to be about 27, 240 nt, which included 38.07 % GC content is in 0349 and 38.13 % GC content in J0304. In general, the genome of HCoV-229E is characterized with a gene order of 5’-replicase ORF1a/b, spike (S), envelope (E), membrane (M), nucleocapsid (N)-3’ [35], Like SARS-CoV-2, the spike protein is the determinant of infections to host cells [36]. The ORF1a/b of HCoV-229E encodes 16 non-structural proteins (NSPs). The *NSP5* encodes the M^pro^ that is required for the replication in the host cells [34]. Given that HCoV-229E is allowed to be studied in BSL2 laboratories, this pathogenic virus is an appropriate model to screen therapeutics for the treatment of both common cold and COVID-19.

Given that the SARS-CoV-2 M^pro^ not only plays a vital role in the cleavage of polyproteins, and there is no human homolog, it is an ideal target for anti-SARS-CoV-2 drug screens and development [37, 38]. It belongs to the family of cysteine proteases and has a Cys-His catalytic dyad, which is an appropriate site to design and screen antiviral drugs [39]. Its high-resolution crystal structure was elucidated in April 2020 [40]. Based on the crystal structure, screening the existing antiviral medicines or designed chemicals revealed that cinanserin, ebselen, GC376, 11a, and 11b showed inhibitory effects on the M^pro^ activity [39–43]. A common feature is that these molecules deliver their carbonyl group (aldehyde group or ketone group) to the thiol of the 145 cysteine residue to form a covalent linkage, thus inhibit the M^pro^ activity. The potential application of these molecules is still under studies to evaluate their effectiveness and side effects. In addition, we recently found that flavan-3-ol gallates, such as (-)-epigallotechin-3-gallate, (-)-catechin-3-gallate, and (-)-epicatechin-3-gallate, and dimeric procyanidins promisingly inhibited the M^pro^ activity [44]. Docking simulation indicated that their inhibitory activity likely resulted from the formation of hydrogen bonds between these compounds and several amino acids in the binding domain of M^pro^.

Flavonols and dihydroflavonols (Fig. 2) are two main groups of plant flavonoids [45, 46]. Quercetin, kaempferol, and myricetin are three flavonol molecules widely existing in plants. Likewise, dihydroquercetin, dihydrokaempferol, and dihydromyricetin are three dihydroflavonol molecules in plants [47, 48]. In general, flavonols and dihydroflavonols are strong antioxidants with multiple benefits to human health [49–58]. Furthermore, studies have reported that quercetin and its derivatives have antiviral activity [59–61]. Based on these previous findings, we hypothesized that flavonols and dihydroflavonols might inhibit the M^pro^ activity of SARS-CoV-2 and HCoV-229E. In this study, to test this hypothesis, we performed docking simulation for three dihydroflavonols, three flavonols, and two glycosylated quercetins. Then, we tested these compound’s inhibition against the recombinant M^pro^ activity of SARS-CoV-2 *in vitro*. More importantly, five available compounds were evaluated to determine their inhibitive activity against the replication of HCoV-229E in Huh-7 cells. The resulting data showed eight compounds effectively inhibited the M^pro^ activity of SARS-CoV-2 and five tested compounds inhibited the replication of HCoV-229E in Huh-7 cells.

**Figure 2.**
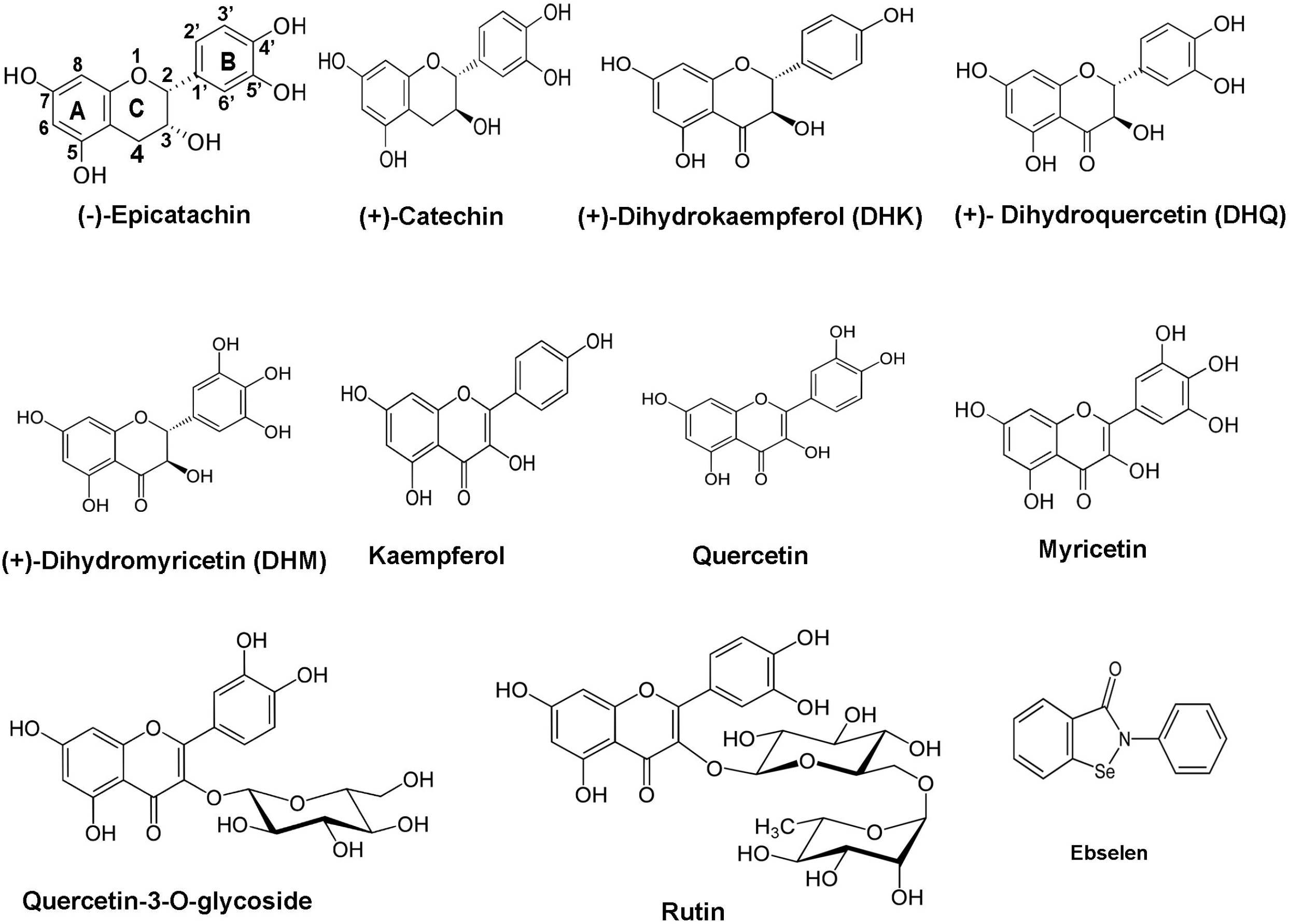
Structures of ebselen and 10 flavonoids. Two flavan-3-ols: (-)-epicatechin and (+)-catechin; three dihydroflavonol aglycones: (+)-dihydroquercetin, (+)-dihydrokaempferol, and (+)-dihydroquercetin; three flavonols aglycones, kaempferol, quercetin, and myricetin; two glycosylated flavonols: quercetin-3-O-glycoside (isoquercitrin), and rutin.

## 2. Materials and methods

### 2.1 Dihydroflavonols, flavonols, cell line, and coronavirus

Flavonols used in this study included kaempferol, quercetin, myricetin, quercetin-3-O-glycoside, and rutin. Dihydroflavonols used were (+)-taxifolin (dihydroquercetin, DHQ), (+)-dihydrokaempferol (DHK), and (+)-dihydromyricetin (DHM). Two flavan-3-ols, (-)-epicatechin and (+)-catechin, were used as compound controls. The ebselen was used as a positive control. These compounds were purchased from Sigma-Aldrich (https://www.sigmaaldrich.com/)

Huh-7 cells, a human hepatocellular carcinoma cell line, are an appropriate to study the replication of different viruses [62–66]. This cell line was used for infection and propagation of virus and for testing antiviral activity of compounds. Human coronavirus 229E (HCoV-229E) is a positive sense and single-stranded RNA virus that infects the human respiratory system [67–71]. HCoV-229E was propagated on Huh-7 cells and tittered by TCID50 assay.

### 2.2 Docking simulation of the SARS-CoV-2 M^pro^

We recently reported the docking simulation of flavan-3-ols, such as epicatechin and catechin (Fig. 2) [44]. Herein, we used the same steps for docking simulation of flavonols and dihydroflavonols in this experiment. In brief, three main steps were completed, protein preparation, ligand preparation, and protein-ligand docking. The first step was protein preparation. The SARS-CoV-2 M^pro^ was used as a receptor to test ligands. Its ID is PDB ID: 6LU7 at Protein Data bank (https://www.rcsb.org/), from which its 3D structure was downloaded to a desktop computer and then was prepared as a receptor of ligand via the Dock Prep tool of UCSF-Chimera (https://www.cgl.ucsf.edu/chimera/). Because M^pro^ contains the inhibitor peptide N3, we removed N3 prior to docking simulation. Hydrogens and charges were added and optimized to allow determining the histidine protonation state. The second step was ligand preparation. The 3D structures of compounds (Fig. 2) were obtained from PubChem (https://pubchem.ncbi.nlm.nih.gov/) and then used as ligands. All structures were minimized by using the minimize structure tool of UCSF-Chimera. Hydrogens and charges were added to the ligands, which were then saved as mol2 format for the protein-ligand docking simulation. The third step was protein-ligand docking. The modeling of protein-ligand docking was performed via the publically available AutoDock Vina (http://vina.scripps.edu/) software. The protein and ligand files were loaded to the AutoDock Vina through the UCSF-Chimera surface binding analysis tools. A working box was created to contain the whole receptor. The box center was set at x =-27, y = 13, and z = 58. The box size was set as x = 50, y =55, and z = 50, which framed the entire receptor to allow free position changes and ligand binding to the receptor at any potential positions.

### 2.3 Docking simulation of the HCoV-229E M^pro^

HCoV-229E needs a M^pro^ (3C-like protease) for its replication in the host cells [72–74]. The M^pro^ is also a target for screening anti-HCoV-229E medicines [75–77]. The sequence of the HCoV-229E M^pro^ was obtained from the GenBank and then used for an alignment and docking simulation. The steps of simulation were the same as described above.

### 2.4 Inhibition assay of the SARS-CoV-2 M^pro^ activity

(+)-DHQ, (+)-DHK, (+)-DHM, quercetin, kaempferol, myricetin, quercetin-3-O-glycoside, rutin, (-)-epicatechin, and (+)-catechin were dissolved in DMSO to prepare a 1.0 M solution. A SARS-CoV-2 Assay Kit (BPS bioscience, https://bpsbioscience.com/) was used to test the inhibitory activity of these compounds. The steps of *in vitro* assay followed the manufacturer’s protocol as performed in our recent report [44]. In brief, each reaction was carried out in a 25 µl volume in 384-well plates. Each reaction solution contains 150 ng recombinant M^pro^ (6 ng/µl), 1 mM DDT, 50 µM fluorogenic substrate, and one compound (0, 0.02, 0.05, 0.1, 0.5, 1, 5, 10, 50, 100, 150, and 200 μM) in pH 8.0 mM Tris-HCl and 5 µM EDTA buffer. GC376 (50 μM) was used as a positive control, while (-)-epicatechin and (+)-catechin were used as two negative controls. The reaction mixtures were incubated for 2 hrs at room temperature. The fluorescence intensity of each reaction was measured and recorded on a microtiter plate-reading fluorimeter (BioTek’s Synergy H4 Plate Reader for detect fluorescent and luminescent signals). The excitation wavelength was 360 nm and the detection emission wavelength was 460 nm. Each concentration of every compound was tested five times. A mean value was calculated with five individual replicates. Plots were built with the percentiles of catalysis versus log [µM] values of concentrations to show the effect of each compound on the M^pro^ activity. Statistical evolution is described below.

### 2.5 Inhibition assay of Human coronavirus 229E

Huh-7 cells were grown in Dulbecco’s Modified Eagle Medium (DMEM) supplemented with 10% fetal bovine serum (10% FBS) and 1% antibiotics. HCoV-229E was propagated on Huh-7 cells. Virus containing supernatants were harvested 72h post infection and stored at −80°C. The virus titer was determined by the Median Tissue Culture Infectious Dose 50 (TCID50) assay in Huh-7 cells.

Then, we performed virus inhibition assays. Huh-7 cells were seeded in 96 well plates at a density of 25,000 cells/well and incubated overnight. HCoV 229-E was diluted in MEM with 1% FBS, 1% HEPES buffer, and 1% antibiotic solution (MEM 1+1+1). The cells were inoculated with HCoV-229E at an MOI of 1 in a total volume of 50 µl. The infected plates were incubated at 35°C with 5% CO2 for one hour. Phytochemicals dissolved in DMSO were added in cell culture medium to the following concentrations: 0 µM, 2.5 µM, 5 µM, 10 µM, 20 µM, and 50 µM.

After one hour, virus and medium were removed from the infected cells and washed once with 200ul of PBS. 100 µl of each compound master mix was added to triplicate wells for each concentration. Virus was allowed to grow in the presence of each compound at 35 °C and 5% CO2 for 24 hours. Supernatants were harvested and virus titers on Huh-7 cells were determined by TCID50 assay [78]. Plates were incubated at 37°C and 5% CO2 for 96 hours, inspected visually for cytopathic effect (CPE) and TCID50/mL was calculated using the Spearman-Kaerber method [79, 80]. A mean value was calculated using three replicates. Plots were built with TCID50/mL versus concentrations to show the effect of each compound on the replication of virus in Huh-7 cells. The minimum level of detection in this assay was 632 TCID50/ml.

### 2.6 Statistical evaluation

One-way analysis of variance (ANOVA) was performed to evaluate the statistical significance. The P-value less than 0.05 means significant differences.

## 3. Results

### 3.1 Ligand-receptor docking of flavonols and dihydroflavonols to the M^pro^ of SARS-CoV-2

Docking simulation was completed with the UCSF-Chimera and AutoDock Vina software to evaluate the binding abilities of flavonols and dihydroflavonols to the SARS-CoV-2 M^pro^. The M^pro^ structure is featured with a substrate-binding pocket (Fig. 3). When the 3D structure of the protein was downloaded from the public database, the peptide inhibitor N3 was shown to bind to this pocket. During protein preparation, N3 was removed for docking. The simulation results showed that (+)-DHQ, (+)-DHK, (+)-DHM, quercetin, kaempferol, myricetin, quercetin-3-O-glycoside, rutin, (-)-epicatechin, (+)-catechin, and ebselen bound to the binding pocket (Fig. 3). The resulting affinity scores for (+)-DHQ, (+)-DHK, (+)-DHM, quercetin, kaempferol, myricetin, quercetin-3-O-glycoside, and rutin ranged from −8.8 to −7.4, lower and better than the score of ebselen (−6.6) (Table 1). The scores among the aglycones of (-)-epicatechin, (+)-catechin, three dihydroflavonols, and three flavonol aglycones were close, either −7.4 or −7.5. These data suggested that dihydroflavonols, flavonols, and glycosylated flavonols could potentially inhibit the M^pro^ activity.

**Figure 3.**
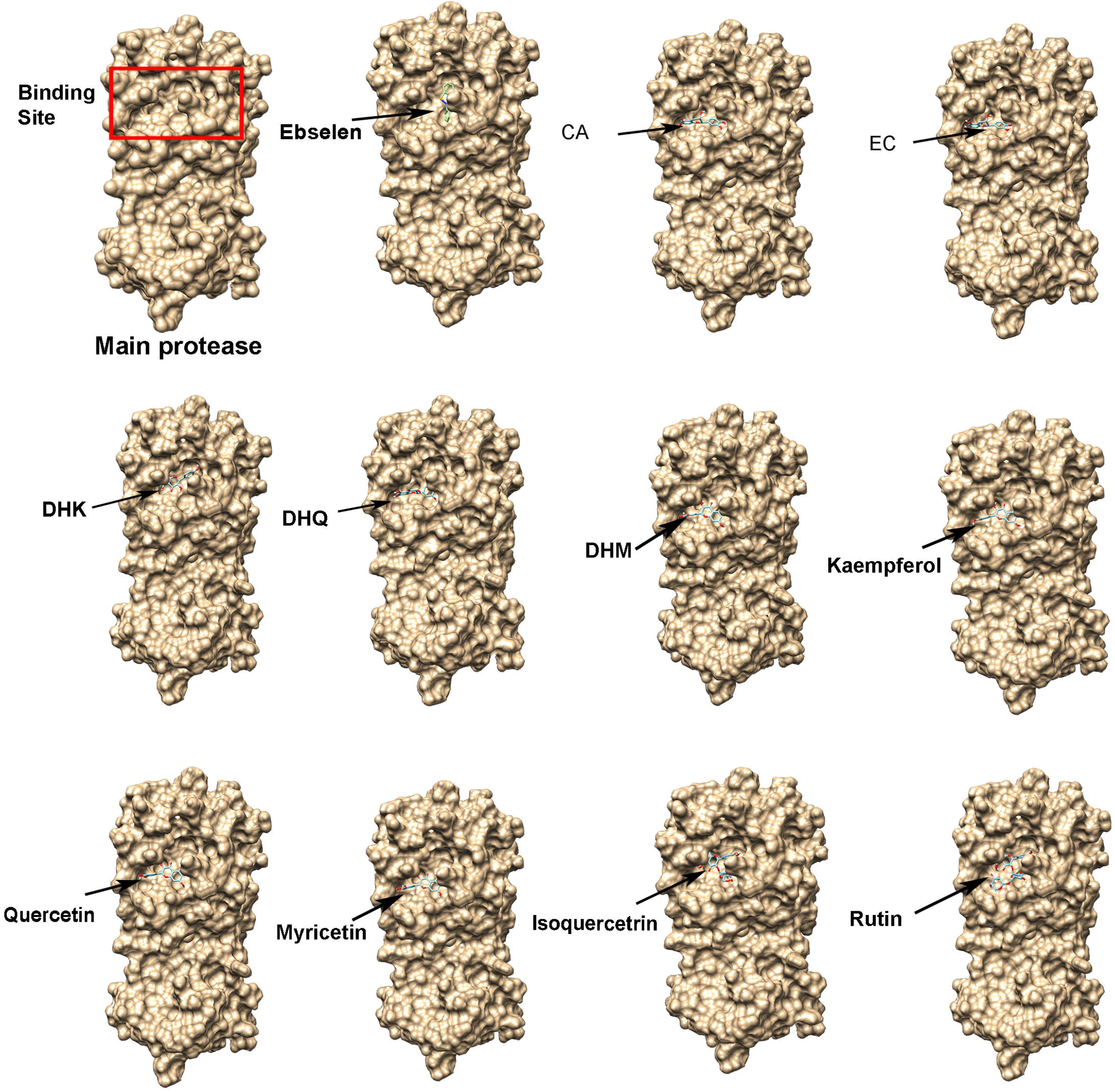
Ligand-receptor docking modeling showing the binding of eleven compounds to the substrate pocket of the SARS-CoV-2 main protease (M^pro^). The first image shows the 3D surface view of the SARS-CoV-2 M^pro^, on which the red rectangular frame indicates the substrate-binding pocket. Eleven flavonoids and ebselen bind to this pocket. Two flavan-3-ols: (+)-catechin (CA) and (-)-epicatechin (EC); three dihydroflavonol aglycones: (+)-dihydroquercetin (DHQ), (+)-dihydrokaempferol (DHK), and (+)-dihydroquercetin (DHM); three flavonols aglycones, kaempferol, quercetin, and myricetin; two glycosylated flavonols: quercetin-3-O-glycoside (isoquercitrin), and rutin.

**Table 1.** Affinity scores of 11 compounds binding to the main proteases of SARS-CoV-2 and HuCoV-229E

| Compounds | Affinity score<br>(SARS-CoV-2) | Affinity score (229E) | Molecular weight<br>(Da) |
| --- | --- | --- | --- |
| Rutin | -8.8 | -7.8 | 610.5 |
| Isoquercitrin | -8.7 | -7.5 | 464.1 |
| Kaempferol | -7.7 | -7.6 | 286.2 |
| DHK | -7.6 | -7.6 | 288.2 |
| DHM | -7.5 | -7.6 | 320.2 |
| Myricetin | -7.4 | -7.1 | 318.2 |
| Quercetin | -7.4 | -7.7 | 302.2 |
| (+)-Taxifolin | -7.4 | -7.8 | 304.2 |
| (+)-catechin | -7.5 | -7.6 | 290.2 |
| (-)-epicatechin | -7.5 | -7.6 | 290.2 |
| Ebselen | -6.6 | -6.0 | 274.2 |

### 3.2 Docking features at the binding pocket of the SARS-CoV-2 M^pro^

As we reported recently [44], the M^pro^ substrate-binding pocket includes four subsites, S1’, S1, S2, and S4. Cys145 is a critical residue located at the space among subsites S1, S1’, and S2 (Fig. 4a) [39, 40]. Several studies have reported that the thoil of the Cys145 residue is crucial for the catalytic activity of M^pro^ and if a compound binds to this residue, it can inhibit the M^pro^ activity [21,39,42]. When ebselen was used as our positive compound for simulation, as we reported recently [44], it bound to this residue featured by three rings facing to the S1 and S1’ subsites (Fig. 4 b). The docking simulation results showed that three dihydroflavonols, three flavonols aglycones, and two glycosylated flavonols bound to 2 to 4 subsites via the Cys145 residue. In three dihydroflavonols tested, (+)-DHK and (+)-DHQ showed a difference in their occupation in the binding site. The A and B rings of (+)-DHK and (+)-DHQ dwelled in the S1’ and S2 subsites and their heterocycle C ring resided in the space between the S1 and S2 subsites (Fig. 4 c-d). The A and B rings of DHM occupied the S1 and S4 subsites and the heterocycle C ring resided in the space between the S1 and S2 subsites (Fig. 4 e). In three flavonol aglycones tested, the occupation of kaempferol was different from that of quercetin and myricetin. The A-ring, B-ring, and heterocycle C-ring of kaempferol resided in the S1’, S2, and the space between S1 and S2, respectively (Fig. 4f). The A-ring, B-ring, and the heterocycle C-ring of quercetin and myricetin dwelled in the S1, S4, and the space between S1 and S2 (Fig. 4 g-h). In comparison, the residing positions of isoquercitrin and rutin were more complicated. The A-ring, B-ring, heterocycle C-ring, and 3-glucose of isoquercitrin occupied the S2, S1’, the space between S1 and S2, and S1 (Fig. 2 i). The A-ring, B-ring, heterocycle C-ring, 6-β-glucopyranose, and 1-L-α-rhamnopyranose of rutin occupied S4, S1’, the space between S1/S2, S1, and S4 (Fig. 4 j). These occupations in the binding sites suggested that these compounds might have an inhibitive activity against M^pro^.

**Figure 4.**
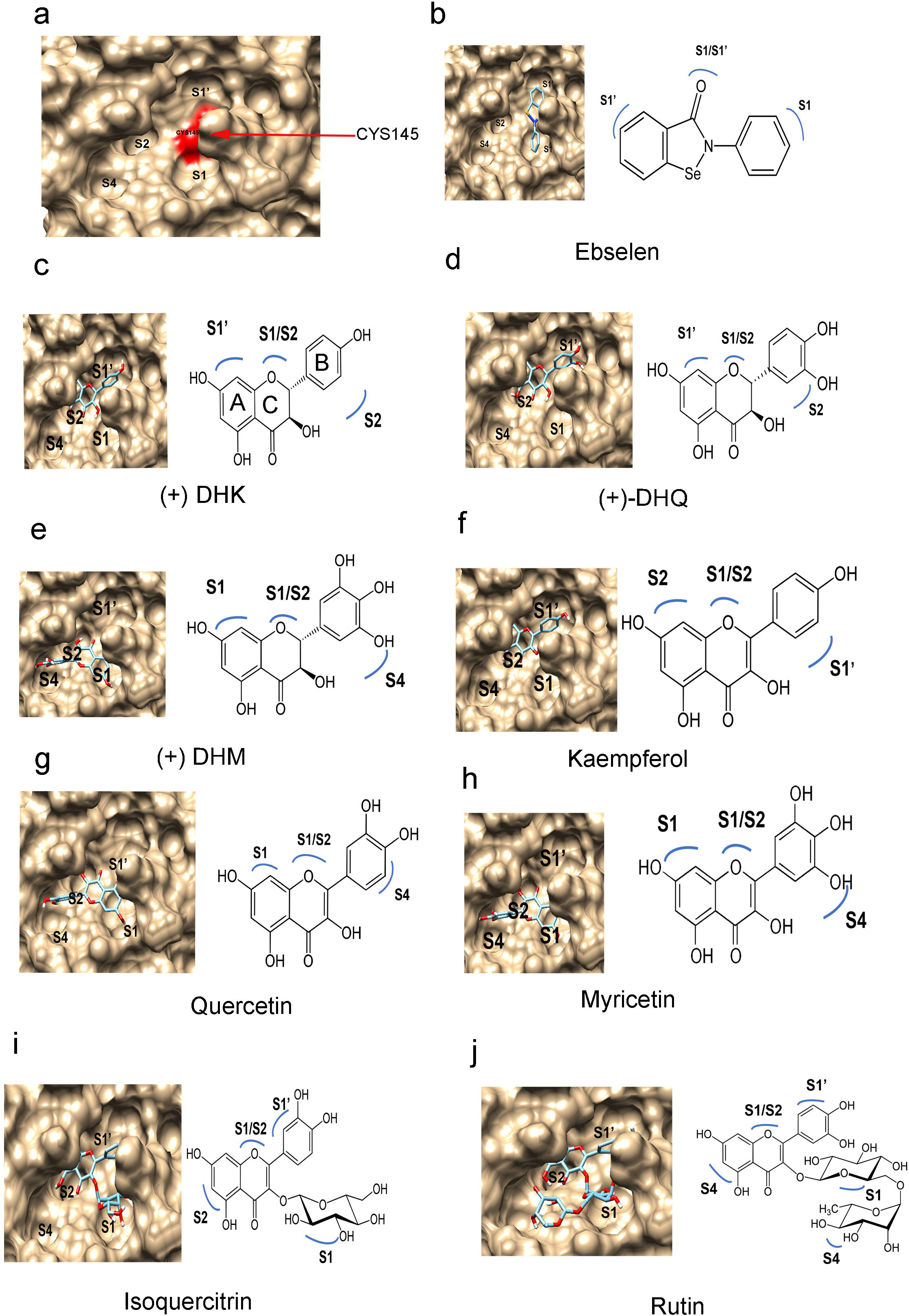
Orientation features of compounds binding to subsites. a, a surface image shows the four subsites in the binding pocket. b-j, images show the binding positions of nine compounds. Two flavan-3-ols: (+)-catechin (CA) and (-)-epicatechin (EC); three dihydroflavonol aglycones: (+)-dihydroquercetin (DHQ), (+)-dihydrokaempferol (DHK), and (+)-dihydroquercetin (DHM); three flavonols aglycones, kaempferol, quercetin, and myricetin; two glycosylated flavonols: quercetin-3-O-glycoside (isoquercitrin), and rutin.

### 3.3 Ligand-receptor docking of flavonols and dihydroflavonols to the M^pro^ of HCoV-229E

The M^pro^ of HCoV-229E was also used for docking simulation. A sequence alignment revealed that the identity of between the two M^pro^ homologs of HCoV-229E and SARS-CoV-2 was 42.81% (Figure 5 a). The binding domains were highly conserved. Furthermore, a 3D modeling revealed that the conformation and binding pocket of the HCoV-229E M^pro^ were similar to those of SARS-CoV-2 M^pro^ (Fig. 5 b and c). The simulation results were the same as those of the M^pro^ of SARS-CoV-2 described above. (+)-DHQ, (+)-DHK, (+)-DHM, quercetin, kaempferol, myricetin, quercetin-3-O-glycoside, rutin, (-)-epicatechin, (+)-catechin, and ebselen could bind to the binding pocket of the HCoV-229E M^pro^ (Fig. 6). The affinity scores of these compounds ranged from −7.8 to −7.1 (Table 1). The scores of rutin and isoquercitrin (two glycosides) binding to the HCoV-229E M^pro^ were −7.8 and −7.5, higher than −8,8 and −8.7, the scores of the two compounds binding to the SARS-CoV-2 M^pro^ (Table 1). This result indicates that compared with the affinity score of quercetin, these two types of glycosylation reduce the affinity scores binding to the SARS-CoV-2 M^pro^, but do not affect the affinity scores binding to the HCoV-229E M^pro^.

**Figure 5.**
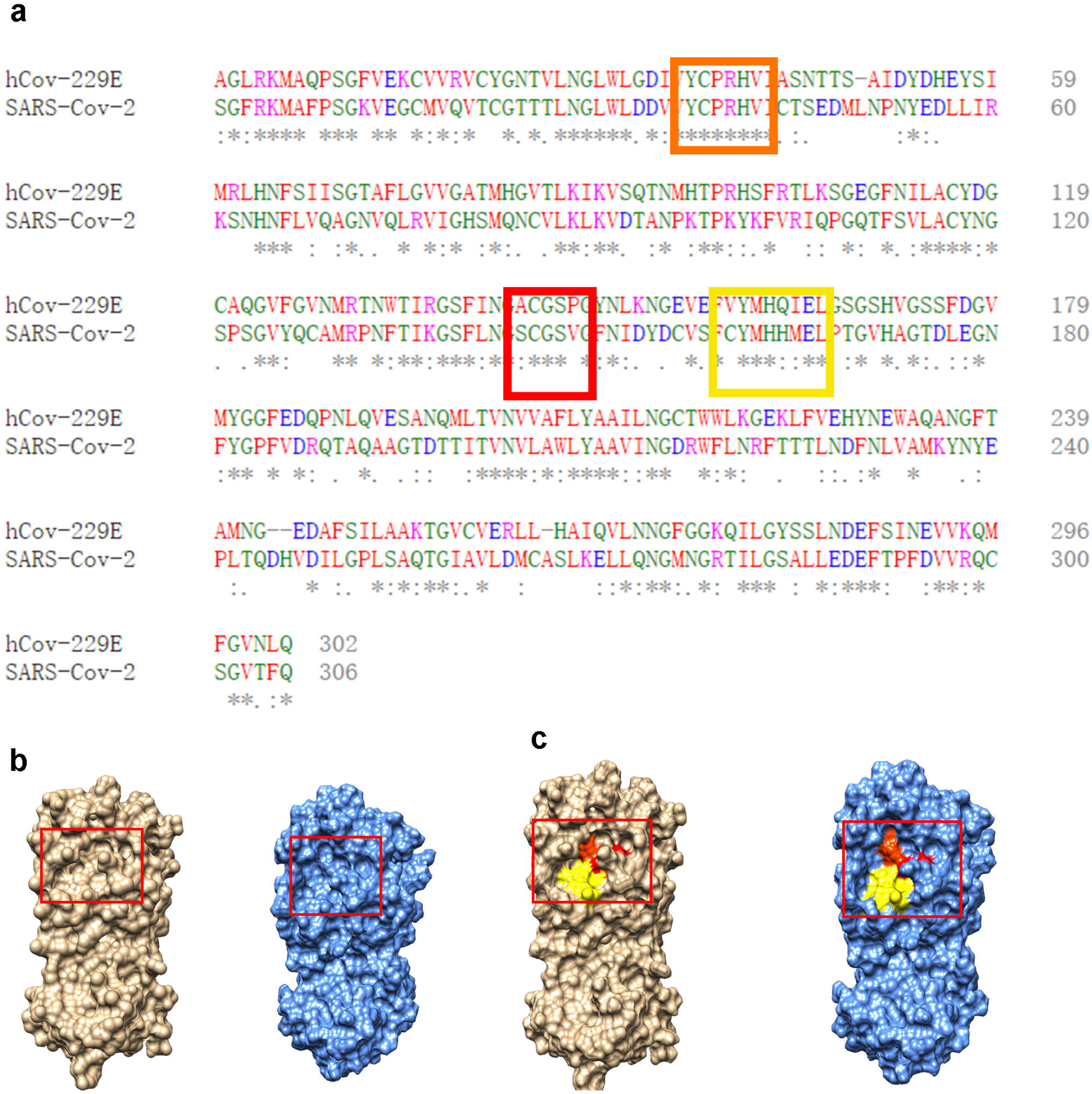
Amino acid sequence alignment of the SARS-CoV-2’s and HCoV-229E’s M^pro^ homologs and comparison of their three dimensional (3D) models. a, amino sequence alignment, in which three rectangle frames highlight three conserved domains forming the substrate binding pocket; b, a comparison of the 3D models of the SARS-CoV-2’s (bronze color) and HCoV-229E’s (blue color) M^pro^ homologs; c, yellowish, orange, and reddish colors showing the binding pocket formed from three conserved binding domains highlighted with three rectangle frames in a, in which the reddish and yellowish spaces include Cys-His catalytic dyad.

**Figure 6.**
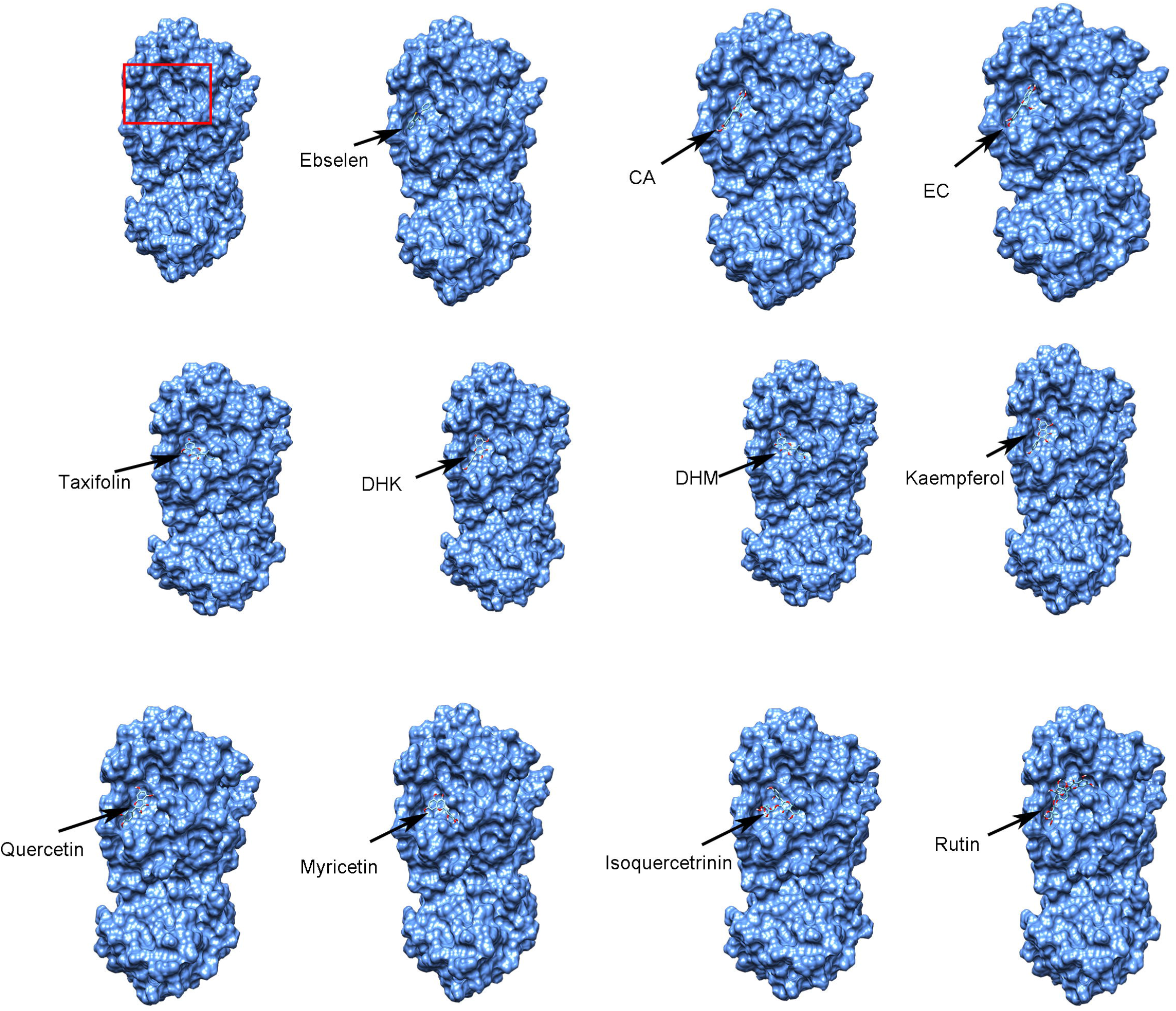
Ligand-receptor docking modeling showing the binding of eleven compounds to the substrate pocket of the HCoV-229E main protease. The first image shows the 3D surface view of the HCoV-229E M^pro^, on which the red rectangular frame indicates the substrate-binding pocket. Eleven flavonoids and ebselen bind to this pocket. Two flavan-3-ols: (+)-catechin (CA) and (-)-epicatechin (EC); three dihydroflavonol aglycones: (+)-dihydroquercetin (DHQ), (+)-dihydrokaempferol (DHK), and (+)-dihydroquercetin (DHM); three flavonols aglycones, kaempferol, quercetin, and myricetin; two glycosylated flavonols: quercetin-3-O-glycoside (isoquercitrin), and rutin.

### 3.4 *In vitro* inhibitory effects of five flavonols and two dihydroflavonols on the SARS-CoV-2 M^pro^ activity

(+)-DHQ, (+)-DHM, quercetin, kaempferol, myricetin, isoquercetin (quercetin-3-O-glycoside), and rutin were used to test their inhibitory effects on the M^pro^ activity. In addition, based on our recent report [44], (-)-epicatechin and (+)-catechin were used as negative controls. The resulting data showed that (+)-DHQ, (+)-DHK, (+)-DHM, quercetin, kaempferol, myricetin, isoquercetin (quercetin-3-O-glycoside), and rutin inhibited the SARS-CoV-2 M^pro^ activity. The half maximum inhibitory concentrations (IC50) were 0.125-12.94 µM (Fig. 7). Among the tested seven compounds, rutin had the lowest IC5 value with the most effectiveness to inhibit the M^pro^ activity (Fig. 7 g), while (+)-DHQ had the highest IC50 value with the lowest inhibitive activity (Fig. 7 d). One hundred µM was used to further compare the inhibitive effects of these compounds on the M^pro^ activity in a given time. The resulting data showed the most effectiveness of rutin (Fig. 7 h). In addition, as we reported previously, (+)-catechin and (-)-epicatechin did not show an inhibitory effect on the M^pro^ activity in the range of concentrations from 0-200 µM. For example, the two compounds did not inhibit the catalytic activity of the M^pro^ at 100 µM (Fig. 7 h).

**Figure 7.**
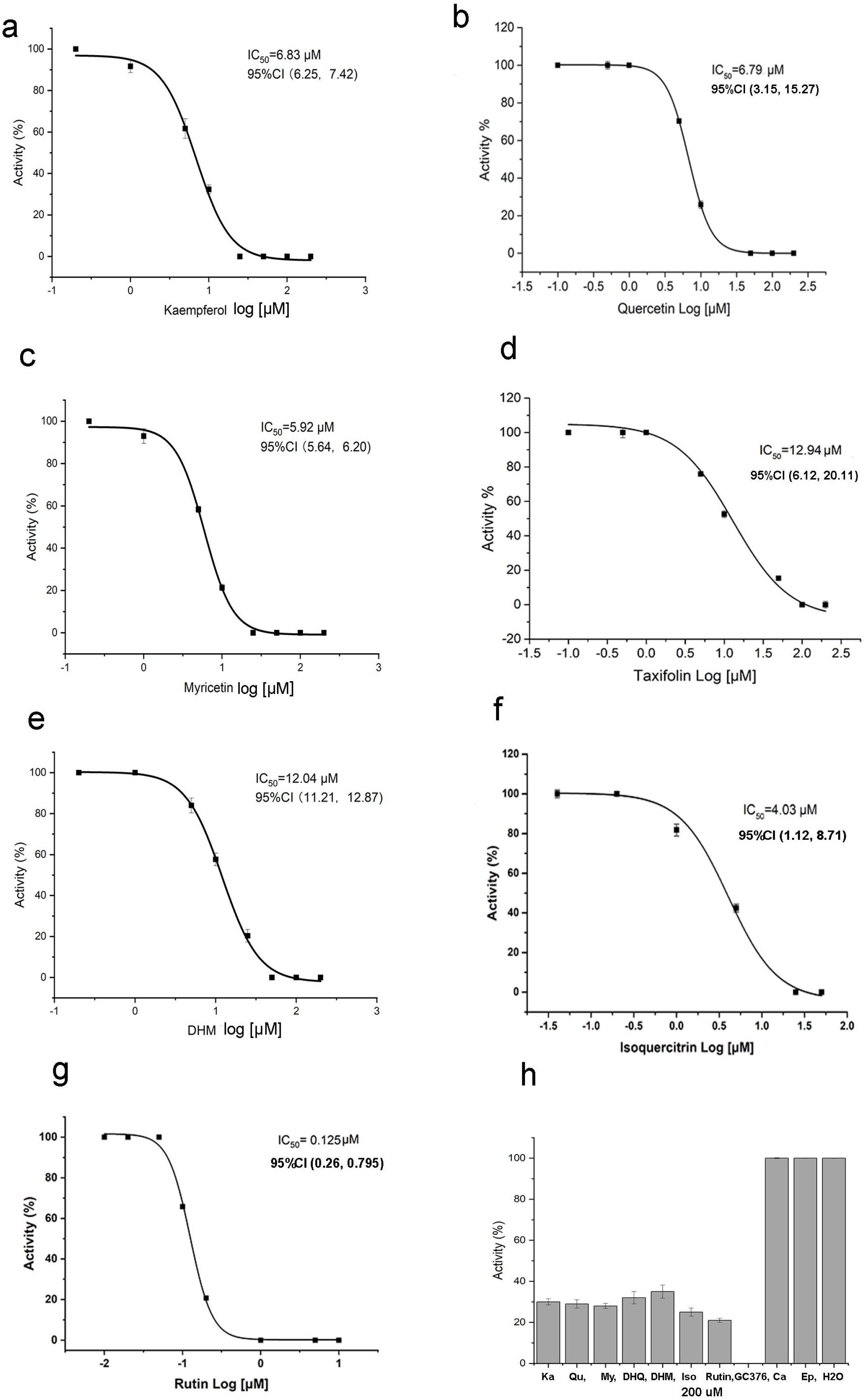
Inhibitory effects of nine compounds on the M^pro^ activity of SARS-CoV-2. a-g, seven plots show the inhibitory curves of seven compounds against the M^pro^ activity. All dots in each plot are an average value calculated from five replicates. IC50 value for each compound is inserted in each plot. “95% Cl” means 95% confidence internal. “(value 1, value 2)” means values in the range with 95% Cl. h, a comparison shows the inhibitory effects of 11 compounds at 100 µM on the M^pro^ activity. GC376 is an inhibitor used as positive control. (+)-catechin, (-)-epicatechin, and water are used as negative controls. Two flavan-3-ols: (+)-catechin (Ca) and (-)-epicatechin (Ep); three dihydroflavonol aglycones: (+)-dihydroquercetin (DHQ), (+)-dihydrokaempferol (DHK), and (+)-dihydroquercetin (DHM); three flavonols aglycones, kaempferol (Ka), quercetin (Qu), and myricetin (My); two glycosylated flavonols: quercetin-3-O-glycoside (isoquercitrin, Iso), and rutin.

### 3.5 Inhibitory effects of five compounds on the replication of HCoV-229E in Huh-7 cells

Quercetin, isoquercetin, taxifolin, were tested their inhibitory effects on the replication of HCoV-229E in Huh-7 cells. In addition, epigallocatechin gallate (EGCG), and epicatechin two examples of flavan-3-ols, were tested. The reason was that we recently reported that EGCG effectively inhibited the SARS-CoV-2 M^pro^ activity, while epicatechin could not [44], however whether they could inhibit coronavirus replication in the host cells was untested. It was essential to test them. The resulting data indicated that all five compounds showed an inhibition against the replication of HCoV-229E in Huh-7 cells (Figure 8). Based on TCID50/ml values, taxifolin started to show its inhibition at 2.5 µM and its inhibitory activity increased as its concentration was increased. Quercetin started to have inhibition at 5 µM. As its concentrations were increased, its inhibitive activities were more effective. At a concentration tested higher 10 µM, quercetin could strongly inhibit the replication of the virus. Its EC50 value was estimated to be 4.88 µM (Figure 8 b). Isoquercitrin strongly inhibited the replication starting with 2.5 µM. EGCG started to show its inhibition against the replication of the virus at 2.5 µM and its inhibition became stronger as its concentrations were increased. It was interesting that epicatechin could strongly inhibit the replication starting at 20 µM.

**Figure 8.**
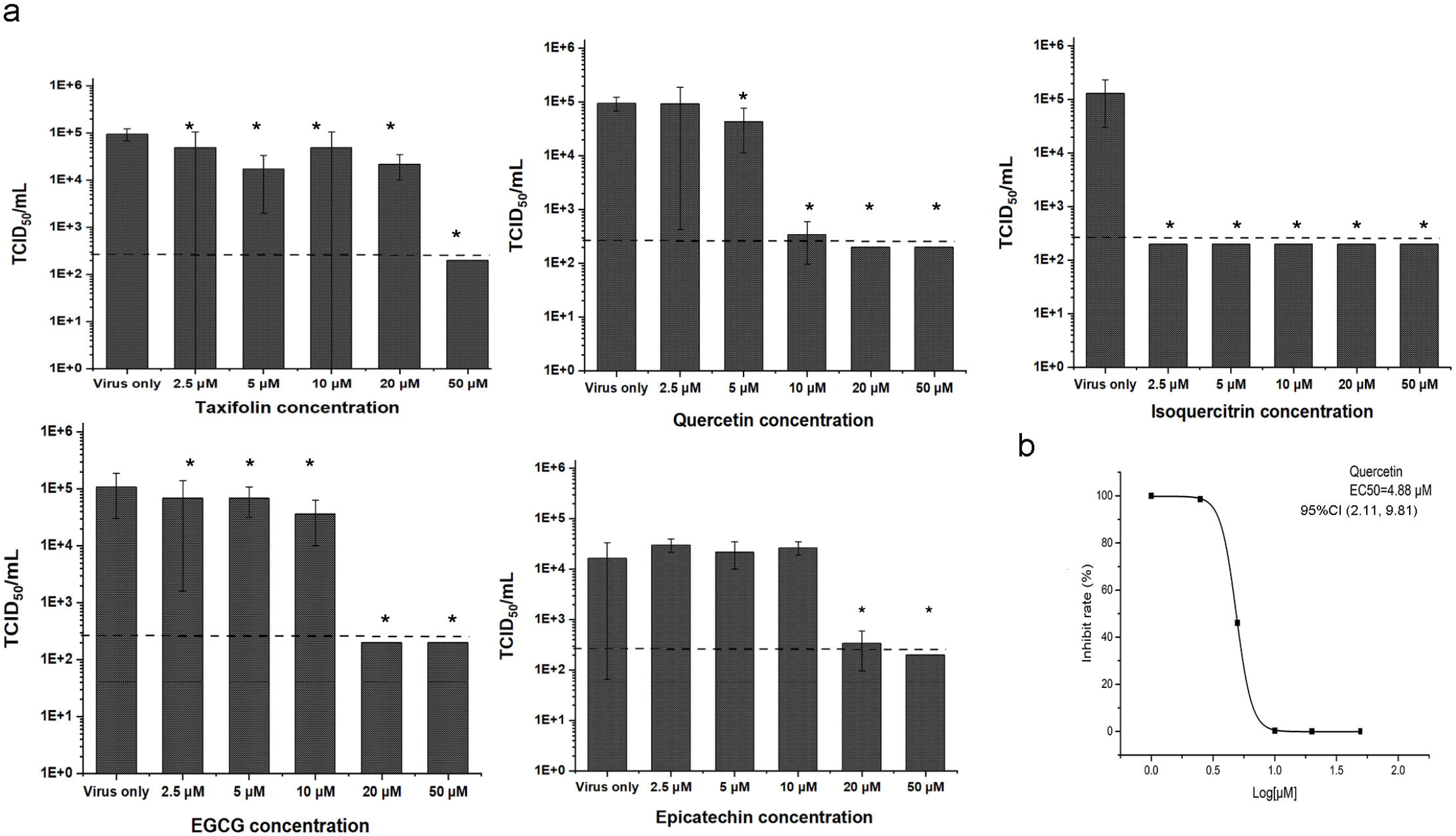
Inhibition of five compounds on the replication of HCoV-229E in Huh-7 cells. a, plots were built with TCID50/mL versus concentrations of each compound. b, this plot was built with the inhibition rate (%) versus log [µM] values to estimate the EC50 of quercetin. “95% Cl” means 95% confidence internal. “(value 1, value 2)” means values in the range with 95% Cl. Bars labeled with “*” means were significant difference compared with control without adding compounds (P-value less than 0.05).

## 4. Discussion

The development of medicines is necessary to complement the use of vaccines to control COVID-19. The SARS-CoV-2 M^pro^ is one of the targets to screen, repurpose, or develop drugs to treat or prevent SARS-CoV-2 [21,22,39]. One strategy is to inhibit the M^pro^ activity via delivering a compound to the Cys145 residue at the space across the region of S1’ and S1 subsites [39]. Ebselen is a small molecule candidate that has been found to inhibit the M^pro^ activity with an IC_50_ 0.46 µM [40]. Its structure featured with three rings was revealed to be an effective vessel to deliver its carbonyl group to the CYS145 residue (Fig. 4 b). We recently reported another strategy. We have found that epicatechin gallate, epigallocatechin gallate, gallocatechin gallate, catechin gallate, and procyanidin B2 could effectively inhibited the activity of M^pro^ via the formation of hydrogen bonds with different amino acids in the binding pocket [44]. Our findings indicated that the formation of peptide bonds was effective to screen more flavonoids to intervene COVID-19. Quercetin and other flavonols are common nutraceuticals with antiviral activities, such as influenza virus, hepatitis B virus, Zika virus, and Ebola viruses [59,81–83]. In this study, we took advantage of our recent strategy to perform this docking simulation of flavonols and dihydroflavonols. These two groups of compounds (Figs. 2 and 4) have C4 keto and 3-OH structures in the heterocycle C-ring. Like flavan-3-ol gallates, the structures of these two groups might have a potential to reside in the space S1 and S2 subsites. In the present study, our ligand-docking simulation showed that these two groups of compounds could bind to the substrate-binding pocket of M^pro^ and occupied their heterocyclic C ring in the crossing region between S1 and S2. Furthermore, the docking results predicted the A-ring and B-ring of two, three, two, and one compounds could bind to S1’ and S2, S1 and S4, S2 and S1’, and S4 and S1’, respectively (Fig. 4). The docking results further showed that a glycosylation of quercetin increased the dwelling capacity in the binding site. Rutin was predicted to occupy all four subsites (Fig. 4 j). The increase of binding subsites was also reflected by the affinity scores of M^pro^-ligands. Rutin and isoquercitrin had the lowest and second lowest score values (Table 1). These data indicated that not only might these compounds have an inhibitive activity but also a lower and better affinity score might indicate a strong inhibition against the M^pro^ activity. Further *in vitro* assays substantiated the prediction of docking simulation. Seven available compounds inhibited the activity of M^pro^ with IC5 values from 0.125 to 12.9 µM. These data imply that these compounds might be potential therapeutics.

Given that SARS-CoV-2 can be only handled in the BSL-3 laboratories, we cannot access this deadly virus to test the effects of these compounds on its replication in host cells. Instead, we selected the less pathogenic HCoV-229E to test the inhibitory activity of these compounds. We hypothesized that inhibitory compounds screened against this virus might be appropriate for the potential therapy of COVID-19. The reason is that like SARS-CoV-2, the replication of HCoV-229E also depends on its M^pro^ activity in human cells and the active site is highly conserved between HCoV229E and SARS-CoV-2. Accordingly, the resulting data might help design medicines for the therapy of both COVID-19 and HCoV-229E respiratory diseases. An amino sequence alignment revealed that the identity of HCoV-229E and SARS-CoV-2 M^pro^ homologs was approximately 48%. The binding domain of substrates between the two homologs was conserved. Docking simulation and the resulting affinity scores further indicated that these compounds could reside in the binding pocket to potentially inhibit the activity of the M^pro^ of HCoV-229E (Fig. 4 and Table 1). Based on these data, we could test five compounds with HCoV-229E. In tested concentrations, taxifolin and isoquercitrin starting from 2.5 µM showed a significant inhibition of HCoV-229E replication in Huh-7 cells. Quercetin could slightly reduce the replication of HCoV-229E at 2.5 µM and significantly inhibited the replication of this virus at higher concentrations (Fig. 8 a). These positive results not only supported the docking simulation results that these compounds bound to the M^pro^ of HCoV-229E (Table 1), but also substantiated the results of *in vitro* assays that these compounds effectively inhibited the SARS-CoV-2 M^pro^ activity (Fig. 7). We previously demonstrated that EGCG could effectively inhibit the activity of the SARS-CoV-2 M^pro^ [44]. Herein, we used it as a positive control. The resulting data showed that EGCG starting with 2.5 µM could significantly inhibit the replication of HCoV-229E in Huh-7 cells.

This positive control result further supported that taxifolin, isoquercitrin and quercetin inhibited the replication of HCoV-29E via the reduction of the M^pro^ activity. In addition, we tested epicatechin, which was not shown to have an inhibitive activity against the SARS-CoV-2 M^pro^ in our vitro assays. It was interesting that epicatechin starting with 20 µM tested could inhibit the replication of HCoV-229E (Fig. 8 a), which was supported by the results of the docking simulation and its affinity score (Fig. 6 and Table 1). Accordingly, this datum indicates the difference between HCoV-229E and SARS-CoV-2. Taken together, these data indicate that HCoV-229E is appropriate substitute to screen inhibitors of SARS-CoV-2 by targeting the M^pro^ of these two viruses.

Quercetin, isoquercitrin and rutin are three common supplements, given that their nutritional values benefit human health [84–89]. Our data suggest that quercetin, isoquercitrin, and rutin might be helpful to intervene COVID-19. These compounds are plant natural flavonoids that their bio-availability, metabolism, and toxicity have been studied extensively [46]. In general, these compounds are safe nutrients sold as supplements or in food products such as onion and common dinner table fruits [90–95]. More importantly, quercetin can be absorbed into the human body from the intestines. A large number of human health studies have reported the presence of quercetin and its derivatives in the blood plasma and their nutritional benefits after consumption [96–99]. For example, the quercetin concentration in plasma was reported to reach 5.0±1.0 µM after the intake of 150 mg in one hour [100, 101]. In addition, these compounds are potent antioxidants [102, 103]. The intake of quercetin can inhibit the oxidation of LDL and prevent the cardiovascular diseases [101, 104] [105]. Moreover, quercetin and its derivatives have strong anti-inflammation activity [106–110]. All of these functions can benefit people’s health.

## Notes

### Competing Interest Statement

The authors have declared no competing interest.

